# Population genomics and pathotypic evaluation of the bacterial leaf blight pathogen of rice reveals rapid evolutionary dynamics of a plant pathogen

**DOI:** 10.1101/704221

**Authors:** Jinshui Zheng, Zhiwei Song, Dehong Zheng, Huifeng Hu, Hongxia Liu, Yancun Zhao, Ming Sun, Lifang Ruan, Fengquan Liu

## Abstract

*Xanthomonas oryzae* pv. *oryzae* (*Xoo*) causes bacterial blight disease, which reduces crop yield by up to 50% in rice production. Despite its substantial threat on food production worldwide, knowledge about its population structure, virulence diversity and the relationship between them is limited. We used whole-genome sequencing to explore the diversity and evolution of *Xoo* during the past 30 years in the main rice-planting areas of China. Six separate lineages were revealed by phylogenomic analysis, with CX-5 and CX-6 predominating in the population for decades. The recent sporadic outbreaks were respectively caused by *Xoo* derived from these lineages especially the two major ones. The lineage and sub-lineage distribution of isolates strongly correlated to their geographical origin, which was found to be mainly determined by the planting of the two major rice subspecies, *indica* and *japonica*. Large-scale virulence testing was conducted to evaluate the diversity of pathogenicity for *Xoo.* We found rapid virulence dynamics against rice, and its determinant factors including genetic background of *Xoo*, rice resistance genes and the planting environment of rice. Genetic background was investigated deeply by comparative genomics, which indicates that transposition events contributing the most to evolution of the *Xoo* genome and the rapid diversification of virulence. This study provided a good model to understand the evolution and dynamics of plant pathogens in the context of interaction with their hosts which are influenced by both geographical conditions and farming practices.

## Introduction

Population genomics is a powerful tool for understanding the formation and evolution of bacterial pathogens of human and some important domestic animals [1–6]. It has also provided new strategies for the detection, prevention and control of important human pathogens [7, 8]. Though plant pathogens are serious threat to food security worldwide, their evolution and dynamics at population scale in the content of modern agricultural practice have not been well understood. A few studies have used genome-based methods to study their population structure, which have provided some new insights into evolution and transmission of some important crop pathogens [9–13].

*Xanthomonas oryzae* pv. *oryzae* (*Xoo*) causes bacterial blight (BB) disease of rice (*Oryza sativa*) and has been considered one of the top 10 plant bacterial pathogens based on its scientific/economic importance [14]. It is a notorious destructive pathogen which can cause a considerable reduction in rice production in both temperate and tropical regions, especially in Asia [15]. On average, BB can lead to 20%∼30% overall yield reduction with some severe cases even causing up to 50% loss [16]. As the world’s largest producer and consumer of rice, China has suffered serious BB outbreaks and huge food loss since the 1930s, with the most serious damage occurring between 1950s and 1980s [17, 18]. Breeding resistant rice cultivars has been proved to be the most effective method for controlling this pathogen. Since 1980s, BB-resistant rice varieties have been widely planted in China and have significantly reduced the loss caused by BB [19–21]. However, sporadic outbreaks still occurred in different areas of China after the year of 2000 [22, 23]. It is unknown whether the recent outbreaks relate to changes in the genetics and virulence of the past *Xoo* populations in China, allowing it to overcome host defenses. Only few studies focused on population structure of *Xoo* from limited rice-planting areas of China were conducted by DNA fingerprinting methods [24–26]. So very little is known about the evolution, spread and dynamics of *Xoo* involved in different outbreaks during the past decades around the whole country in the context of rapid development of modern agricultural science and technology.

The virulence differentiation among different strains of *Xoo* which was evaluated by defining different races or pathotypes, was frequently studied to infer its virulence dynamics and diversity against rice host. Strains of the same race share a common pathogenic phenotype in a set of tested host cultivars. Rice lines carrying different resistance genes (R genes) determine the race of *Xoo*. Up to now, tens of races of *Xoo* were identified in different rice-planting locations worldwide [22,27–31]. The pathotypes of *Xoo* showed huge diversity among different locations and rapid dynamics along the isolated times. It was predicted that many factors including genetic background of *Xoo* contributing to its virulence diversity [28–30]. However, many crucial details were overlooked due to the intrinsic low sensitivity and stability of the typing method.

Different resistant cultivars carrying bacterial blight R genes have been planted in the different Asian countries for decades, including China [32, 33]. Resistance from R genes carried by rice cultivars plays a crucial role of determining *Xoo* evolution during its interaction with rice hosts. Recent study has showed that R gene *Xa4* has largely affected the race composition of *Xoo* from the Philippines [33]. The pathogenic diversity of *Xoo* in China has been studied previously [22,23,25,26]. Race compositions showed huge diversity and rapid dynamics among different areas along different time points [34, 35]. However, it is unknown the genetic basis for the pathogenicity diversity and dynamics of *Xoo* from China, which means the relationship between genetic population and the ability *Xoo* to overcome different R genes has not been studied. Moreover, the evolutionary force behind the genetic and pathogenic diversity and dynamics is also not clear.

In this study, a comprehensive population genomic study was combined with a large-scale determination of virulence of *Xoo* that have been isolated in the past 30 years in China, to obtain a framework of genetic dynamics and virulence diversity over time and space. Results from this work shed new light on the evolution of this important plant pathogen.

## Results

### Population structure of *Xoo* from China

To understand the population structure of *Xoo* in China, genomes of the 247 strains were sequenced, including 237 isolated from China, 4 and 6 representative ones from Japan and Philippines, respectively (Table S1). Reads were mapped to the genome of PXO99^A^; SNPs were called by Samtools. After the prophage region, repeat sequences and recombinant regions being filtered, the core SNPs contained 5,271 variable sites. Two methods were used to outline the population structure based on the core SNPs, Maximum likelihood (ML) phylogenetic analysis and a tree-independent hierarchical Bayesian clustering (BAPS).

All *Xoo* from China were clustered into 6 lineages (CX-1 to CX-6), with more than 70% members belonging to CX-5 and CX-6 (Figure 1). The three major rice production areas, including South China, Yangtze Valley area and North China, were displayed in Figure 2a. We annotated the lineage-specific tree with the time and space information of the *Xoo* strains isolated (Figure 1, 2b and 2c). CX-1 and CX-2 were most frequently represented in South China at the year of 2003, of which most isolates were from Yunnan province. The *Xoo* belonging to CX-3 were isolated from North China in 1984, 2003 and 2014, with most of which from Northeast China in 2014. CX-5 and CX-6 were nationally distributed during the past 30 years; while most members in CX-5 were dominant in South China and Yangtze Valley, and CX-6 were more frequently from North China and Yangtze Valley.

**Figure 1.**
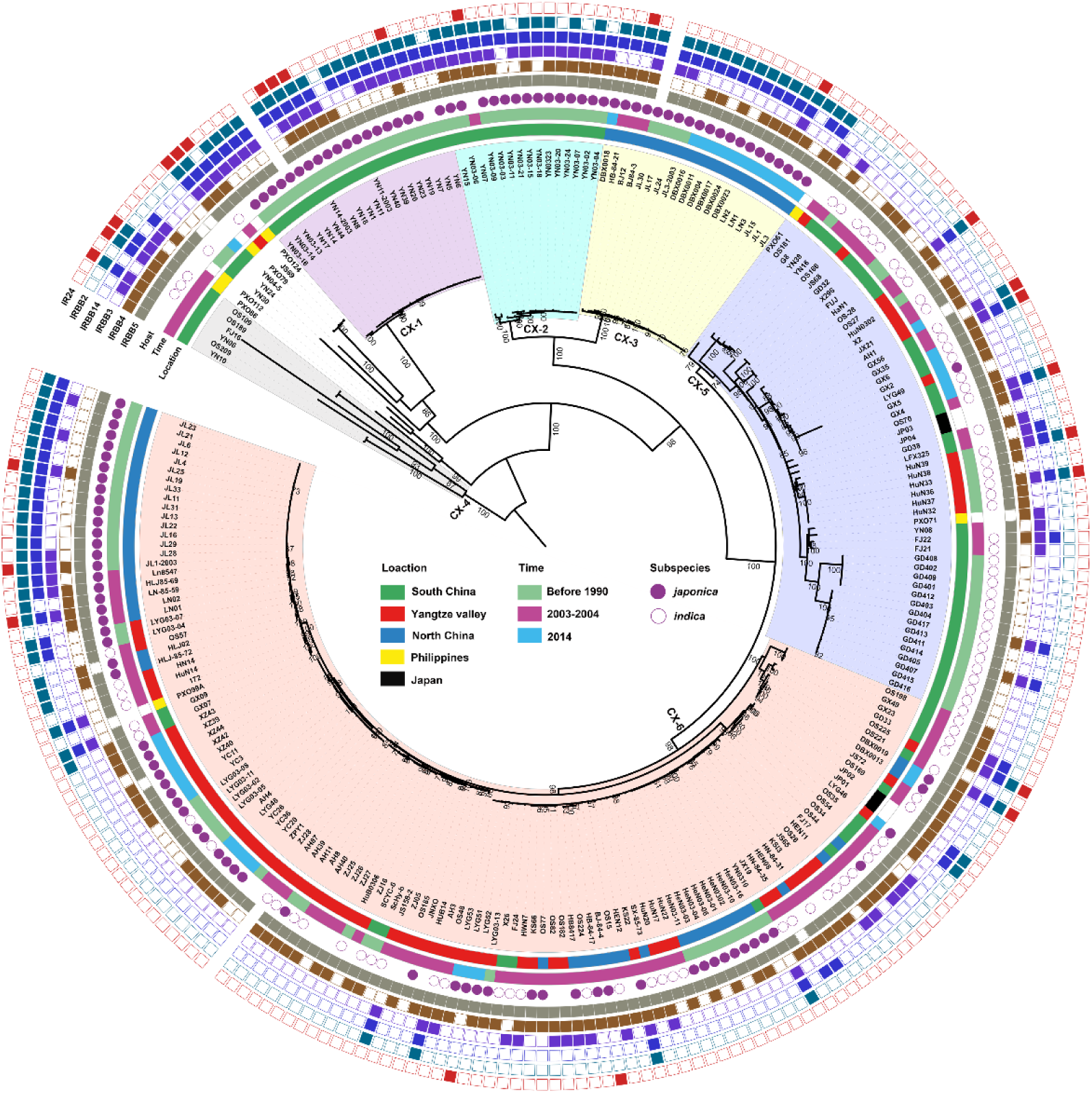
Population structure of *X. oryzae* pv. *oryzae*. The maximum likelihood tree was inferred by RAxML with the generalized time-reversible model and a Gamma distribution to model site-specific rate variation based on the core SNPs of all the *Xoo* genome sequences. The six lineages were decorated with different colors. From inner to outer, the first circle refers to major rice planting areas, the second one describes isolated times of these strains and the third one represents the rice subspecies which the *Xoo* strain isolates. The outmost 6 squares represent the virulence analysis of these *Xoo* against 6 different rice near-isogenic lines with one known resistance gene in each line, with solid square for resistance and hollow one for sensitivity. Bootstrap support values were calculated from 500 replicates, and only values of > 70% were labelled.

**Figure 2.**
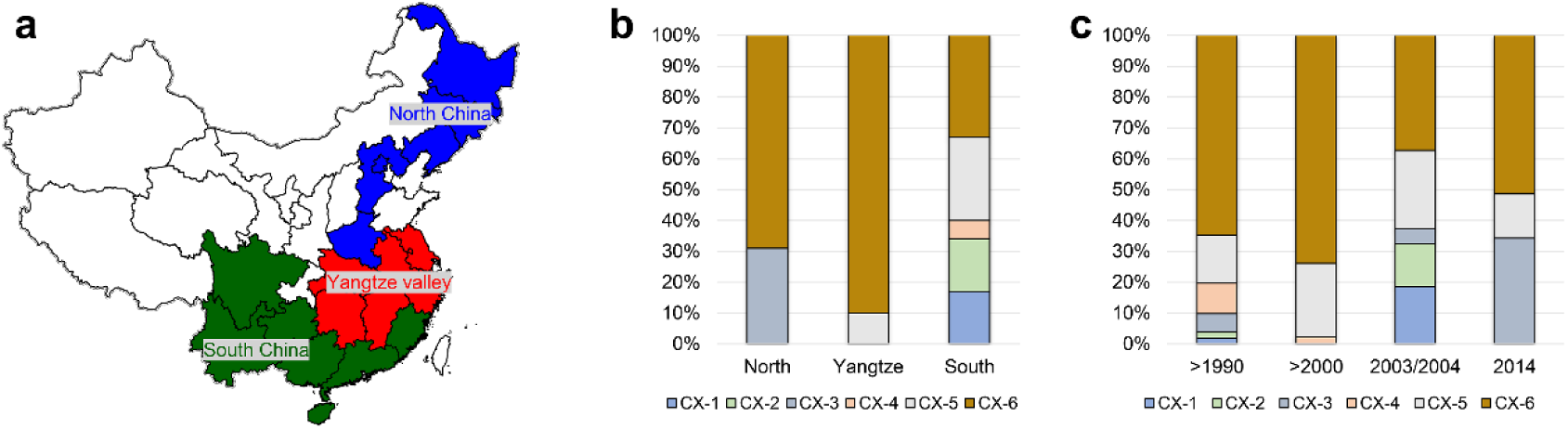
Distribution of the 6 major Chinese *X. oryzae* pv*. oryzae* lineages. Pathotypes of *Xoo* were determined on the following 13 near-isogenic lines, each carrying a specific R gene, IRBB2 (*Xa2*), IRBB3 (*Xa3*), IRBB4 (*Xa4*), IRBB5 (*xa5*), IRBB14 (*Xa14*). IR24 was used as a susceptible check. (a) The map of the 3 major rice producing areas in China. (b) Distribution of the 6 *Xoo* lineages among the 3 rice planting areas. (c) Dynamics of the 6 *Xoo* lineages during the past 30 years.

### Phylogeography of CX-5 and CX-6

To investigate more details about evolution and dynamics of *Xoo* in China, we focused on the two major lineages, CX-5 and CX-6. Phylogenetic analysis based on core SNP alignment (Figures 3a and 3b) and pairwise SNP distance between isolates were performed for both lineages (Figure 3c). Both CX-5 and CX-6 can be divided into 5 sub-lineages, respectively (Figure 3a and 3b).

**Figure 3.**
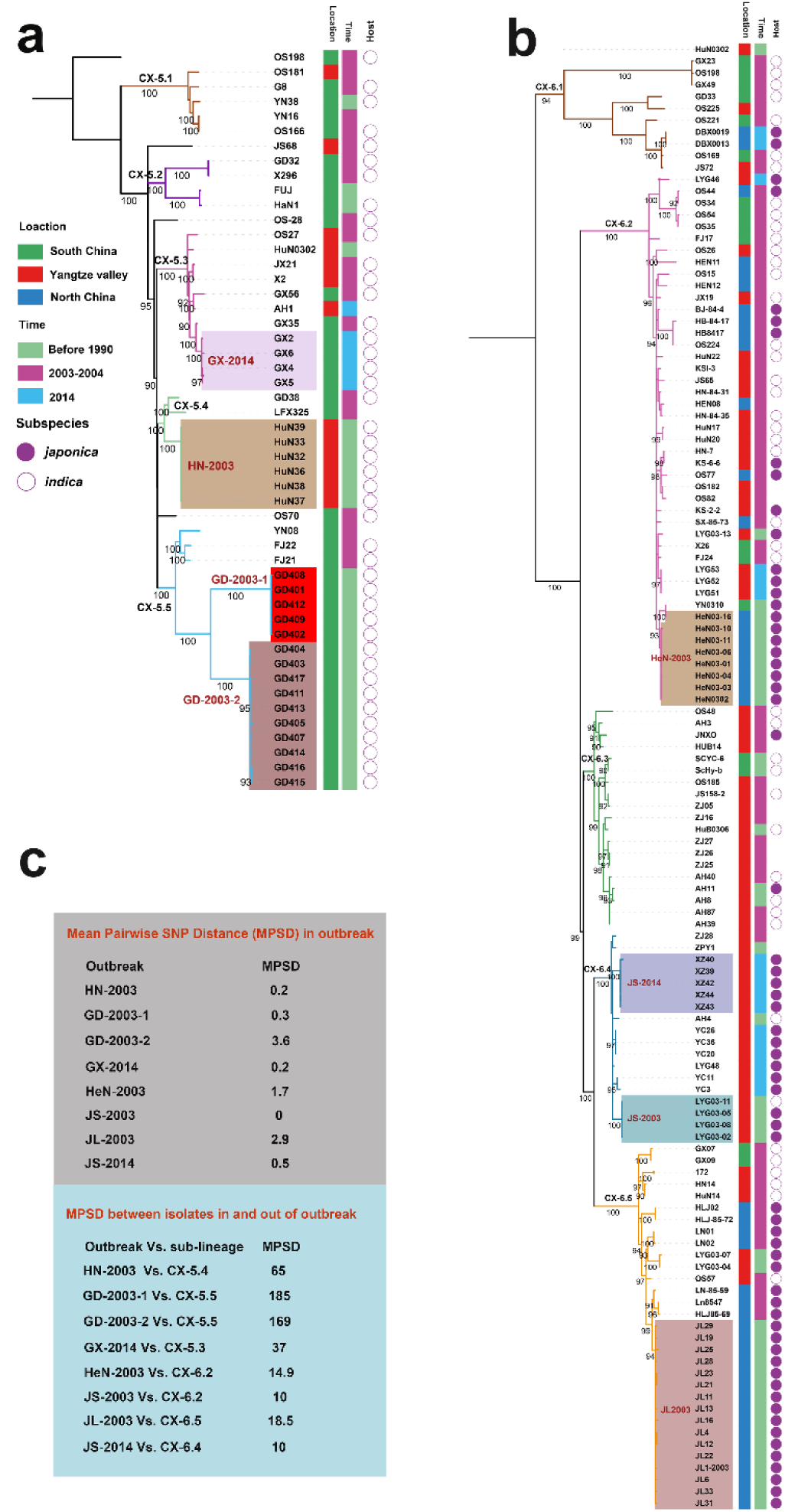
Phylogeography of CX-5 and CX-6. Maximum likelihood phylogeny of Lineage CX-5 (a) and CX-6 (b). Different sub-lineages were displayed by different branch colors. Bootstrap support values were calculated from 500 replicates. The recent outbreaks are represented by different backgrounds. Isolated information of *Xoo* including location, time and rice subspecies were shown in three colored strips. (c) Mean Pairwise SNP Distances (MPSDs) of *Xoo* from the outbreak, and between one in an outbreak and the other one out of the outbreak but in the same sub-lineage were displayed by two different boxes beside the tree.

CX-5.1, CX-5.2, and CX-5.5 were restricted in South China, while CX-5.3 and CX-5.4 can be found in some places of both South China and Yangtze Valley (Figure 3a). When time was considered, we found that all these sub-lineages persisted at these places for at least 30 years. Then we focused on the recent sporadic outbreaks being attributed to this lineage. Besides clustered together on the tree, we considered epidemiologically and genomically linked outbreak with an average SNP pairwise distance less than 5. In 2003, the outbreak in Hunan province of Yangtze Valley was caused by CX-5.4 (HN-2003), while the outbreak in Guangdong province of South China ascribed to two sub-population of CX-5.5 (GD-2003-1 and GD-2003-2). One cluster of CX-5.3 caused outbreak in Guangxi province of South China in 2014 (GX-2014).

Seven out of 10 isolates in CX-6.1 were from different places of South China. This sub-lineage was the most diverse one in CX-6 with long branches. In CX-6.2, *Xoo* from North China and Yangtze Valley were dominant on different branches, respectively. CX-6.3 and CX-6.4 were most frequently found in Yangtze Valley, while *Xoo* from North China predominated in CX-6.5. Similar as CX-5, all these sub-lineages have contributed to BB on rice for a long time in China. These sub-lineages totally caused at less 4 local outbreaks in 2003 and 2014. In 2003, the local outbreaks in two provinces of Yangtze Valley (Henan, and Jiangsu) were caused by CX-6.2 (HeN-2003) and CX-6.4 (JS-2003), respectively, while the one in North China (Jilin province) was ascribed to CX-6.5 (JL-2003). Outbreak in Jiangsu province (Yangtze Valley) in 2014 was caused by one sub-population of CX-6.4 (JS-2014).

### Chinese *Xoo* diversity and dynamics in the context of rice host

There are two major domesticated rice subspecies *Oryza sativa japonica* and *indica* existing in different areas of China; and most of rice varieties were derived from these two subspecies [34, 35]. Rice planted in high-altitude areas of Yunnan province from South China and North China belong to subspecies *japonica* (http://www.ricedata.cn). *Xoo* isolated from these areas were respectively clustered into 3 lineages, CX-1, CX-2, CX-3 and one sub-lineage of CX-6 (CX-6.5) (Figure 1 and Figure 3b). Lineage CX-5 mainly contained *Xoo* isolated from rice in South China where subspecies *indica* is planted (Figure 3a). CX-6 also contained *Xoo* isolated from rice of *japonica* from Yangtze Valley (CX-6.4), *indica* from Yangtze Valley (CX-6.3) and some *indica* from South China (CX-6.1), indicating the widest adaptation to different rice varieties of *Xoo* in this lineage (Figure 3b). In CX-6.2, old *Xoo* were dominant on indica, while recent isolated ones were almost from *japonica*, suggesting a putative host switch in this sub-lineage.

Taken together, the distribution of Chinese *Xoo* lineages appeared to be impaired by biogeography and rapid dynamics over isolated time, which may be mainly due to the distribution and dynamics of the two major subspecies of rice, *japonica* and *indica*.

### Diversity of *Xoo* from different counties of Asia

To place *Xoo* from China into a global context, phylogenomic analysis was performed on the core genome SNPs of 109 *Xoo* genomes available in Genbank (including 100 from India, 8 from the Philippines and 1 from Japan), and the 247 sequenced ones in this study. Phylogenetic analysis revealed a similar topological structure of Asian *Xoo* to that of Chinese *Xoo*, with small differences in some branches (Figure S1). Eleven genetic lineages were identified for the *Xoo* population. *Xoo* from India and the Philippines were respectively assigned into PX-A to PX-C and IX-I to IX-V according to previous studies [36, 37]. Eight lineages displayed strict geographic distribution features, most of them being specific to one or at least a limited amount of rice-planting areas. Besides the three Chinese lineages (CX-1, CX-2 and CX-3), PX-B and PX-C were mainly represented in the Philippines, and IX-I to IX-III were from India. CX-4 contained old *Xoo* from China and some recently isolated ones from India belonging to IX-V. For CX-5 and CX-6, besides both represented the major *Xoo* from China, they also contained a few *Xoo* from Japan, India and Philippines, with PX-A of Philippines being nested within the former one and IX-IV from India being encompassed among the members of the last one, indicating frequently transmission of these 2 lineages not only among different places of China but also among different Asian countries.

### Rapid virulence dynamics of *Xoo* strains isolated in China during the past 30 years

As describe above, the population structure *Xoo* maintains relative stability during the past 30 years. We next inferred the pathogenic diversity and dynamics of these *Xoo* strains. Based on the interactions between *Xoo* strains and rice lines, most of the tested *Xoo* were classified into 9 pathogenic races (Table S1and Figure 4a). The top two races with most members, R5 and R8, contain 26% and 18% of the isolates, respectively.

**Figure 4.**
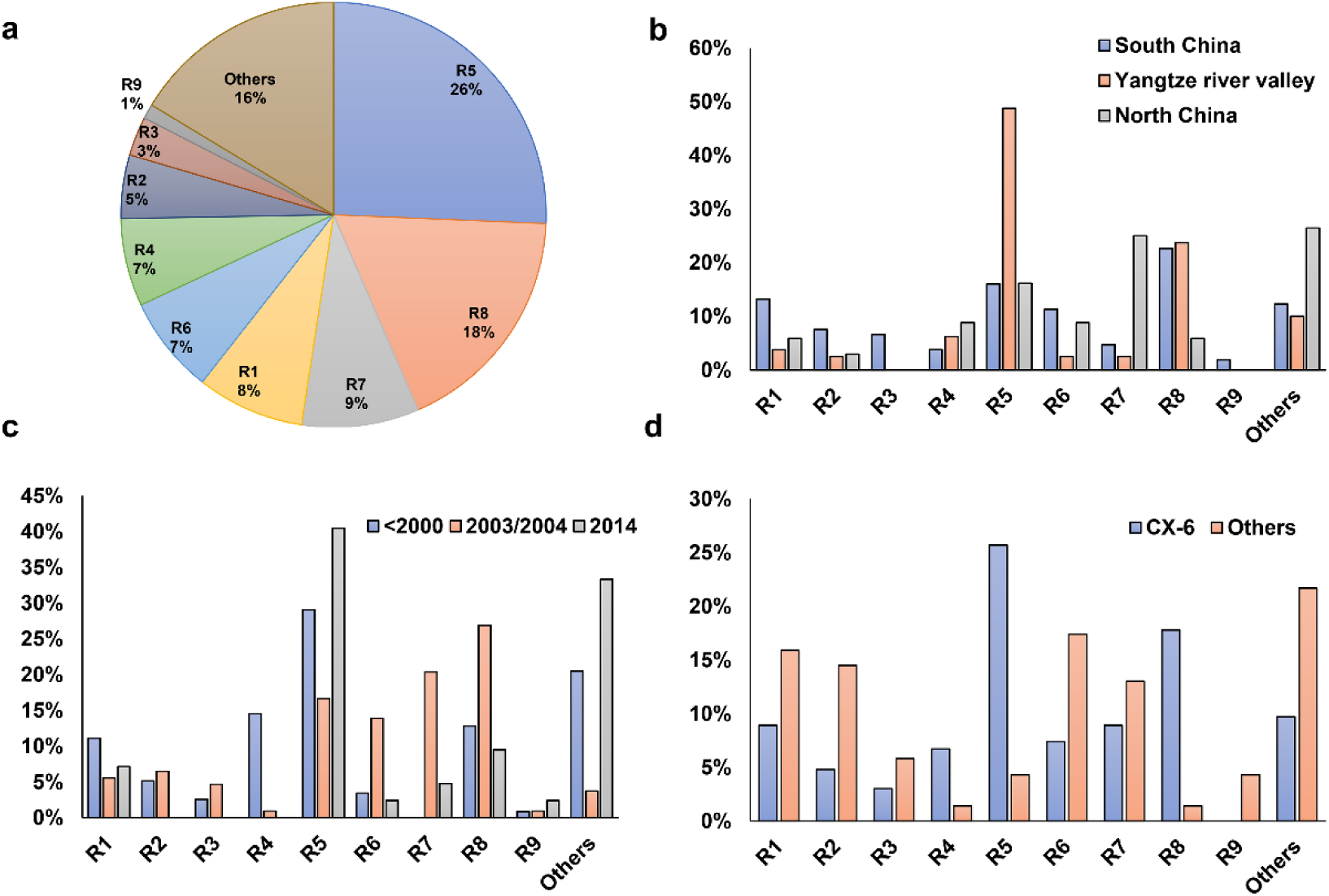
Virulence analysis and race classification of *X. oryzae* pv*. oryzae* China. Pathogenicity profile of each race corresponding to was shown in Table S1. (a) Proportions of the 9 *Xoo* races. (b) Distribution of the 9 races among the 3 major rice planting areas in China. (c) Dynamics of the 9 *Xoo* races during the past 40 years. (d) Distribution of the 9 races in Lineage CX-6 and others.

The three major rice-planting regions had different race compositions (Figure 4b). Yangtze River Valley mainly contained *Xoo* from races R5 and R8, with the respective percentages of the total population being 50% and 30%, respectively. *Xoo* isolates from North China mostly belongs to R7 and R5, together accounts for about two third of the total population. Five of the nine *Xoo* races were well represented in the *Xoo* population from South China, with each of them account for more than 10% of the total population. The virulence dynamics of *Xoo* during the last 30 years was inferred by analyzing race distribution in China at different period (Figure 4c). Strains isolated before the year of 2000 were mainly of the races of R5, R4, R8 and R1, while most *Xoo* isolated in 2003–2004 were R8, R7, R5 and R6; and about 45% of the isolates in 2014 were R5.

We also found different lineages have different race compositions, indicating genetic background playing important roles in the interaction between *Xoo* and rice hosts (Figure 4d). Compared to other lineages, the CX-6 *Xoo* isolates fall mostly within the races of R5, R8 and R4, while have reduced representation in R1, R2 and R6. Moreover, the rapid dynamics of pathogenic features can be inferred by focusing on the races of *Xoo* isolated from the same places at the same time and belonging to the same lineage (Figure 1). For example, most *Xoo* from Yunnan, one province of South China, in 2003 were clustered in CX-1 and CX-2. The 19 isolates in CX-1 represented 8 of the 9 races described above, and the 14 strains in CX-2 belonged to 6 races. Even *Xoo* isolates from the same outbreak were allocated into different races, with 16 *Xoo* in JL-2003 belonging to 3 different races and 10 isolates in GD-2003-2 representing 2 races.

### Resistance gene of rice contributes to the interaction between *Xoo* and rice

As each of the near-isogenic rice lines used in this study contains one known resistance (R) gene, we studied the interaction between *Xoo* and rice host by focusing on the capacity of *Xoo* to overcome the resistances conferred by these genes (Figure 1, Tables 1 and 2). When the time of isolation was considered, we found that *Xoo* from different times have different capacity to overcome different R genes (Table 1). R gene *xa5* showed the most effective resistance against *Xoo* all the time. At different periods, more than 90% *Xoo* could not infect the rice line IRBB5, which contained gene *xa5*. The resistance mediated by R gene *Xa3* was effective during the period of 1990 to 2004. However, rice containing this R gene only showed resistance against 5% of *Xoo* in 2014. Rice cultivars carrying R gene *Xa4* are resistant to more than 60% of *Xoo* isolated from 1970s to 2000 and 2014. However, they are only resistant to about 30% of the isolates recovered in 2003 and 2004.

**Table 1.** Resistance of different rice lines to Xoo strains isolated at different times. Shown is the percentage of *Xoo* strains isolated at different periods that are unable to cause disease.

| **Time** | **IRBB2** | **IRBB3** | **IRBB4** | **IRBB5** | **IRBB14** | **IR24** |
| --- | --- | --- | --- | --- | --- | --- |
| **(# of strains)** | **(*Xa2*)** | **(*Xa3*)** | **(*Xa4*)** | **(*xa5*)** | **(*Xa14*)** | **(*Xa18*\*)** |
| <1990 (35) | 37 | 0 | 97 | 94 | 25 | 20 |
| <2000 (69) | 25 | 58 | 64 | 100 | 28 | 12 |
| 2003/2004 (97) | 49 | 35 | 34 | 100 | 56 | 7 |
| 2014 (42) | 36 | 5 | 79 | 90 | 33 | 5 |

\*: Recurrent parent *Xa18*

**Table 2.** Resistance of different rice lines to Xoo strains of different lineages. Shown is the percentage of *Xoo* strains of different lineages that are unable to cause disease.

| **Lineage**<br>**(# of strains)** | **IRBB2**<br>**(*Xa2*)** | **IRBB3**<br>**(*Xa3*)** | **IRBB4**<br>**(*Xa4*)** | **IRBB5**<br>**(*xa5*)** | **IRBB14**<br>**(*Xa14*)** | **IR24**<br>**(*Xa18*\*)** |
| --- | --- | --- | --- | --- | --- | --- |
| CX-1 (19) | 89 | 79 | 42 | 100 | 95 | 21 |
| CX-2 (15) | 73 | 93 | 80 | 100 | 100 | 7 |
| CX-3 (20) | 100 | 20 | 70 | 100 | 99 | 10 |
| CX-5 (53) | 11 | 32 | 47 | 96 | 17 | 11 |
| CX-6 (123) | 29 | 25 | 62 | 98 | 28 | 6 |

\*: Recurrent parent *Xa18*. CX-4 was not included because of few isolates it contained.

To determine whether the *Xoo* genetic background contributes to the interaction between *Xoo* and rice lines, we studied the resistance capacity of R genes against *Xoo* from different lineages (Table 2). We found that resistance provided by R genes against *Xoo* are indeed depended on the genetic background of *Xoo*. R genes *Xa2* and *Xa14* effectively confer resistance against *Xoo* from Lineages CX-1, CX-2 and CX-3, preventing infection by more than 70% of *Xoo* strains from these lineages. Similarly, R gene *Xa3* and *Xa4* showed potential resistance against *Xoo* from CX-1 and CX-2, and CX-2 and CX-3, respectively. No R gene except *xa5* conferred resistance to more than 50% of *Xoo* strains from CX-5 and CX-6, which is consistent with the situation that CX-5 and CX-6 had much more extensive distribution across different areas of China during the past decades, comparing to the other Chinese *Xoo* lineages. Among the sub-lineages of CX-5 and CX-6, *Xa2* was only effective against *Xoo* of CX-6.5, with effectiveness of 90%, while *Xa4* could confer the resistance against more than 60% *Xoo* from these two lineages except for CX-5.1 and CX-6.5; *Xa14* showed potential medium level of resistance against *Xoo* from CX-6.1 and CX-6.5 (Table S2).

### Genome dynamics of *Xoo* is mainly determined by transposition events

To study the genomic evolution contributing to the rapid dynamics of virulence, several *Xoo* isolates from different lineages were chosen to complete genome for comparative genomics. All proteins of the 22 *Xoo* with complete genomes were clustered into 7542 groups, with 3513 belonging to core genes. The pan genes plot showed that Asian *Xoo* has an open pan genome (**Figure S2**).

It was reported that *Xoo* genome contains numerous insert sequence elements, so we firstly studied the distribution of transposase genes to understand the dynamics of *Xoo* genomes. Each *Xoo* genome contained more than 800 transposase genes, taking up above 16% of the total genes, with most of them from IS5, IS3, IS1595, IS701, IS30 and IS630 (Figure 5 and Table S3). In the core gene sets, 8%, (266 of 3543) were predicted to encode transposases. However, 71% (1251 of 1771) and 62% (942 of 1523) of the distributed gene and unique gene families were transposases (**Figure S3**). It suggests that transposon elements contribute the most to the diversity of *Xoo* genomes. Based on the analysis of gene presence/absence analysis, the core genome of *Xoo* can be split into about 200 conserved blocks by less conserved genome fragments (Figure 5). Among the blocks, there were genome segments with more sequence diversity, and most transposases were in these regions.

**Figure 4.**
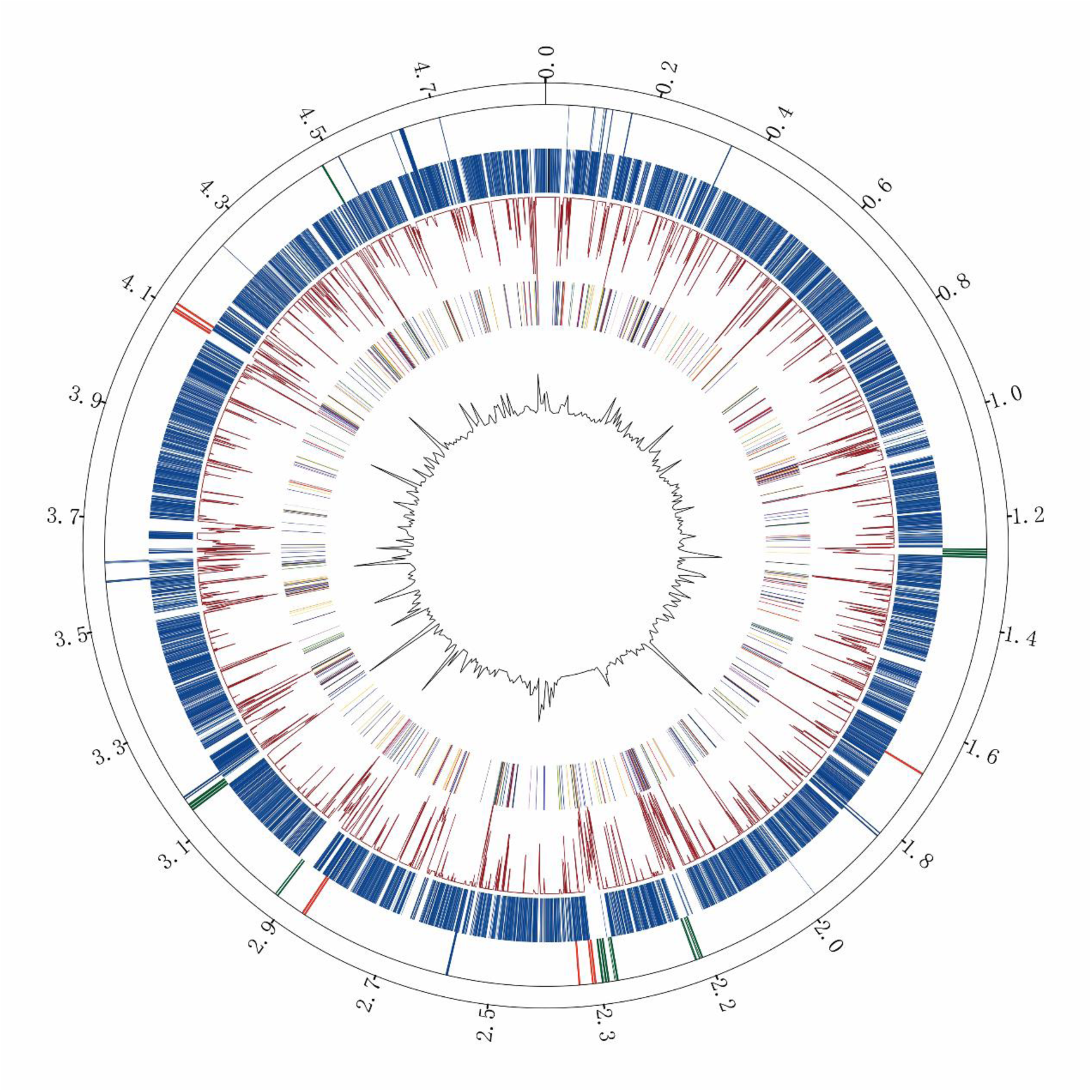
Core and pan genome of *Xoo*. The results showed here is from the comparative analysis of the 22 *Xoo* complete genomes, using OS198 belonging to Lineage CX-6 as reference. From the inner to the outmost, (1) Circle 1, the black break line, SNPs per 10kbp; (2) Circle 2, the colorized lines, different families of transposase genes (magenta for IS3, blue for IS5, yellow for IS30, green for IS630, red for IS701, orange for IS1595, black for others); (3) Circle 3, the red break line, numbers of genome having that gene family (ranging 1 to 22); (4) Circle 4, the dark blue lines, positions of core gene; (5) Circle 5, the colorful lines, presences of type III effector genes, with green for TALEs in OS198, red for TALEs in other Xoo genomes, blue for other T3Es; (6) Circle 6, scales according to genome of OS198.

As a rice pathogen, the host selection was reported to play crucial roles on the evolution of *Xoo*. Type III effectors (T3Es) were found to be crucial to the pathogenicity of *Xanthomas* spp. on plant hosts. Among them, transcription activator-like effectors (TALEs) (Avirulence protein AvrBs3) are the most important in *Xoo* during the interaction with rice. TALEs function as transcriptional factors activate expression of target genes in the host in a sequence specific manner, resulting in enhanced bacterial growth and development of disease symptom (Bogdanove et al 2010). Most TALEs from *Xoo* activate the expression of *SWEET* genes in rice to promote susceptibility [32]. Significant variations in the number of TALEs was observed among the 22 *Xoo* with complete genomes, ranging from 11 to 21 (Table S4). Half of the TALE families (21 out of 42) were only predicted in one or two isolates. And some families only distributed or lost in some special lineages. *Xoo* from the same lineage had some common and different families of TALEs. Even *Xoo* isolated in the same geographical location at the same time with the same phylogenetic relationship, JL28 and JL33, contain different compositions of TALE families. This suggests that interaction among rice and *Xoo* can determine rapid diversity of TALE effectors. When locations of TALE genes were identified, we found that almost all of them were in the variable regions where enrich in transposase genes (Figure 5). Conversely, other T3Es were very conserved, almost all of them located in the conserved genome blocks with less transposons (Figure 5).

## Discussion

Most *Xoo* clusters identified by traditional molecular typing methods from several Asian countries showed strict geographical distribution [28-30,38]. Previous report focusing on *Xoo* from India through population genomics defined 5 lineages, with each of them showing a restricted-region distribution [37]. In this study, based on population genomics, we found that there is a general correspondence between the areas of *Xoo* isolated and most of Asian lineages and some sub-lineages in CX-5 and CX-6 (Figures 1, 3 and S2). Moreover, Asian *Xoo* was reported to be very different from those of Africa and North America regions [39, 40]. This suggests that the geographical factors as affected by climate, soil types, or the choice of different rice cultivars and varieties may influence on the diversity of *Xoo*. Though thousands of rice cultivars have been planted in different areas of China, almost all of them were derived from the two subspecies, *indica* and *japonica*, or hybrid varieties between them (http://www.ricedata.cn). Information of the *Xoo* host subspecies in China indicates that variety of rice was one of the major factors forces on the formation and dynamics of Chinese *Xoo* population. Rice varieties planted in different counties and different areas of a country are determined by geographical environment, climate factors and local agricultural policy [41–43]. Taken together, geography-specific rice cultivars in different countries or different areas in the same country contribute the most to the genetic diversity of *Xoo* infecting them.

Development resistance for novel cultivars of crops and vegetables, is proved to be the most effective strategy fighting against bacterial and fungi pathogens [44–47]. But the durability of plant resistance can be neutralized by evolutionary changes in pathogen populations. Knowledge about pathogen population evolution and dynamics under the force from the resistant plant are crucial for further cultivar development [48]. Though resistance breeding of rice have effectively prevented the unprecedented outbreak of BB since 1990s, sporadic outbreaks occurred in different areas of China after the year 2000 [49]. Genomic epidemiology analysis indicates all *Xoo* strains causing the recent epidemic outbreaks except those in CX-1 were derived from the ones leading to heavy losses in the history. On the other hand, only the minor lineage CX-4 was not represented by the recent outbreaks, so the diversity of the *Xoo* population was not reduced due to wide planting of BB-resistant rice varieties. For the current practice of rice production, the major BB threat in the three rice-planting areas of China is still mainly from CX-5 and CX-6 as they were present in history, which call for different detection and prevention strategies for different areas and rice varieties.

The virulence diversity of *Xoo* in China against rice was well investigated during the past 30 years, however, the relationships between population genetic structure and pathotype of *Xoo* were not explored well [22-24,26,49]. This study shows that races of *Xoo* from South China were more diverse than those from North China and Yangtze Valley areas. In line with this, phylogenetic analysis revealed that *Xoo* from South China are more diverse compared to other places of China (Figure 1). Some rice growing regions in South China, particularly in the Yunnan province, have three harvests per year and use different rice cultivars for each harvest [50, 51]. The greater genetic and virulence diversity of *Xoo* thus could be caused by a greater diversity of rice cultivars in use. Our result also showed that there were diverse races in the same lineage, for example, 85% *Xoo* from lineage CX-6 could be assigned to 8 races (Figure 4d), indicating rapid virulence dynamics in the same lineage. Moreover, the isolates from the same local outbreak even belonged to different races. Although *Xoo* strains isolated in China were consistently assigned to one of the 6 Chinese lineages, the dominating population of *Xoo* races in specific areas displayed rapid dynamics and high level of diversity during the past 30 years (Tables 1 and 2). Rapid virulence dynamics with relative stability of the standing genetic variability of *Xoo* population indicates fast adaptation to host during interaction for this plant pathogen.

In China, since the 1980s, *indica* hybrid rice (especially Shanyou 63) had been planted on a large scale [52]. Later, bacterial blight resistant rice varieties, mainly the *india* rice carrying *Xa4* and *japonica* rice carrying *Xa3*, were grown on a large scale for a long time [31, 52]. Soon after these rice varieties widely used for rice production, the effectiveness of these R genes conferring resistance against infection causing by *Xoo* was significantly compromised (Table 1). Actually, the resistance abilities of these two R genes were also overcome frequently by *Xoo* population in other Asian countries [30,38,53]. R gene *xa5* showed very good resistance (above 95%) to *Xoo* from all lineages. Previously, isolates of *Xoo* having compatibility with *xa5* were only reported from Indian Lineages IX-I and IX-III, and some Philipps and Korea isolates [27,54,55]. The only 2 Chinese *Xoo* from Yunnan province (YN24 and YN04-5) belonging to IX-I and IX-III showed good infection ability to rice with *xa5*, suggesting putative transmission of *Xoo* from India to South China. Besides, several isolates from CX-5 and CX-6, especially those obtained recently, can overcome the resistance contributed by *xa5*, indicating different origins of these resistance compatibilities (Tables 1 and 2). Since rice variants with gene *xa5* were less used in China, the development of its compatibility could not be due to a direct selection pressure of this R gene.

It was reported recently that different *Xoo* population from the Philippines evolved different mechanisms to adapt to the R gene *Xa4* [22]. Chinese *Xoo* Isolates from different lineages and sub-lineages showed distinct capabilities to fight against the same R gene. Overall, lineages and sub-lineages with stricter geographical distribution can be more easily defeated by R genes than the wider distributed ones (Tables 2 and S2). The underlying genetic determinations of these differences call for advanced comparative genomics and wet experiments. However, there is no lineage or sub-lineage with all isolates incompatible with one of the tested R genes, which means that *Xoo* with different genetic backgrounds can adapt to different rice hosts with variable resistances. This suggests that the introduction of new R genes could reduce the loss causing by *Xoo* but can’t change the genetic diversity of the *Xoo* population, as no lineage or sub-lineage can be eliminated thoroughly. This is partially explained by that R genes usually restrict the pathogens to the initial infection site, but have no killing abilities [56].

Most of the R genes cloned up to now were reported to mediate resistance associated with the TALEs during the interaction between rice and *Xoo* [57, 58]. The TALE proteins are the most import effectors used by *Xoo* to interact with rice, enforcing virulence, proliferation, and dissemination against rice [59]. All the TALE genes were found in the several variable regions where enriched in transposase genes belonging to families IS3, IS5, IS30 and IS701, respectively. We also find that other T3Es didn’t have much diversity. It is predicted that transposons containing these kinds of transposons contribute to the rapid dynamics of virulence of *Xoo* isolates mainly through affecting the compositions and structure of TALEs. Similar situation was reported that Tn3-like transposon was proved to play a major role of the spread of pathogenicity determinants in *Xanthomonas* spp. [60]. Actually, transposon-rich genome regions were reported to play crucial roles in rapid evolution of effectors after host changes in the Irish potato famine pathogen *Phytophthora infestans* [61].

## Conclusion

The introduction of several resistance genes into rice tremendously reduced the damage of this notorious plant pathogen; only sporadic events were reported after 2000s [19,21–23]. However, this study found that the genome evolution and virulence dynamics of *Xoo* are extraordinary rapid, indicating its ability to overcome the resistance conferred by the current resistance genes and cause the potential for large-scale outbreaks in the future. It is therefore prudent to continue surveillance of disease outbreaks caused by this plant pathogen, and to develop novel strategies for its control. The rice varieties and resistance genes were predicted to be crucial to the evolution and dynamics of *Xoo*. The subspecies of *Oryza sativa* used for rice production and resistance gene introduction into the rice planted were mainly decided by human. The results from the population genomics and large-scale virulence tests provided new insight into how plant pathogens evolve and spread in the context of human agricultural activities. Resistance genes from rice can be used to fight against *Xoo*, but *Xoo* can rapid develop strategies to overcome these resistances. Transposon elements were proved to play a crucial role in the evolution progress during the interaction between pathogen and it rice host. The interaction between plant pathogen and host was reported to conform to zig-zag-zig model [62], this study presented a well-documented example for co-evolution of a pathogen and its host.

## Materials and Methods

### Genome sequencing

*Xoo* strains from 3 countries were used in this study (Table S1). The major strains were isolated from China. And the represented *Xoo* strains for 4 different races from Japan and 7 different races from Philippines were also sequenced.

The bacterial DNA of overnight cultures (30 °C) was extracted and purified using the Easy-DNA kit (Invitrogen, USA) following the manufacturer’s protocol. Total DNA was sequenced by with Illumina HiSeq 2000, 2500, or 4000 to produce pair-end reads with lengths of 100, 125 and 150 bp. For each *Xoo*, raw reads were assessed with the FastQC tool (https://github.com/s-andrews/FastQC) and quality filtered using Trimmomatic (https://github.com/timflutre/trimmomatic). The filtered reads were error-corrected by library with Quake to produce clean reads [63]. Most of the analysis were conducted with the PGCGAP pipeline developed by our bioinformatic team [64]; and more details are described as below.

### Mapping and SNP calling

To study the population structure of *Xoo* from China, reads from each strain sequenced was mapped onto the reference genome PXO99^A^, which is used as a model strain for studying the interaction between *Xoo* and rice host and is the first *Xoo* with complete genome sequence [65], using BWA [66]. Variant detection was performed using Samtools mpileup (https://github.com/samtools/samtools) and filtered with a minimum mapping quality of 30. SNP was excluded if its coverage was less than 10% or more than 200% of the average coverage, if it was not supported by at least 5 reads on each strand. Phage regions and repetitive sequences of the PXO99^A^ genome were predicted by PHASTER (http://phaster.ca) and RepeatScout (https://bix.ucsd.edu/repeatscout), respectively. Snippy (https://github.com/tseemann/snippy) was used to generate the whole genome alignment of all studied *Xoo*, and then the recombination was detected by Gubbins (https://sanger-pathogens.github.io/gubbins) on the core genome alignment after prophage and repeat regions were filtered. SNPs located within phage regions, repetitive sequences or recombinant regions were excluded.

### Assembly and whole genome alignment

To study the population structure of Asian *Xoo*, genomes from different locations were downloaded from the Genbank as follows: 100 from India, 9 from Philippines, 1 from South Korea and 1 from Japan [36, 37]. We performed genome assembly from all the strains sequenced in this study. For each strain, the pair-end clean reads were assembled using Spades [67]. After a few evaluations, different sets of k-mer values were chosen for Spades according to different read lengths. The final assemblies were obtained by filtering out contigs with few reads supported or with lengths lower than 200 bp (Table S1). Whole genome alignment was carried out on all the genomes obtained from public database and assembled here by Parsnp using the genome of PXO99^A^ as reference [68]. Core SNPs were extracted from the core genome alignment by snp-sites (https://github.com/sanger-pathogens/snp-sites). Recombination of the core genome alignment was inferred by Gubbins. The final SNPs were obtained by filtering out the prophage region and repeat sequences mentioned above, and the recombined regions predicted here.

### Phylogenetic analysis

The SNPs-based maximum likelihood (ML) phylogenies were built by RAxML with the generalized time-reversible model and a Gamma distribution to model site-specific rate variation [69]. Bootstrap support values were calculated from 500 replicates. *Xoo* population structure of China was defined with the hierBAPS module of the BAPS software, which delineates the population structure by nested clustering [70]. Three independent iterations with upper population sizes of 5, 10, and 15 were used to obtain optimal clustering of the population. Phylogenetic trees were annotated and visualized by iTOL (https://itol.embl.de/).

### Comparative genomics of compete genomes

Ten isolates were selected for sequencing by long-read sequencing platform based on their positions on the phylogenetic tree and the races they represented. High quality total DNA of each isolate was sequenced by the PacBio RS II sequencing platform using P6-C4 chemistry to a mean fold coverage of 150. De novo assembly was conducted by HGAP3 [71].

For more comprehensive analysis of the genomic dynamics of *Xoo*, complete *Xoo* genomes available on Genbank were included. All the genomes were annotated by Prokka [72]. The TAL effectors of each genome were identified and classified by AnnoTALE [73]. Pan and core genome analysis was conducted by the tool Roary [74], based the gff format file constructed by Prokka. The inflation value of 2 was used for the MCL clustering.

### Virulence evaluation of *Xoo* strains

Virulence of *Xoo* was assesses by inoculating 6 near-isogenic rice lines, each carrying a specific R gene, IRBB2 (*Xa2*), IRBB3 (*Xa3*), IRBB4 (*Xa4*), IRBB5 (*xa5*), IRBB14 (*Xa14*) and IR24 (*Xa18*). IR24 was used as a susceptible check. Seeds of all lines in this study were obtained from the China National Rice Research Institute (CNRRI). Three weeks after sowing on a seedbed, seedlings were uprooted and transplanted into the field. The management of the rice plants in the field was proceeded as usual. Bacterial strain was cultivated on NA plates at 28 °C for 48 h. Then the bacterial colonies were suspended in sterile ddH_2_O with concentration adjusted to 3×10^8^ cfu/ml before inoculation. The leaf-clipping method was used to test the virulence of all the *Xoo* strains [75]. Fifteen top fully expanded leaves per rice plant were inoculated with each strain. The lesions length was measured after 21 days post inoculation. The ratio of the lesion length compared to the whole leaf length was calculated as described previously [76]. When the ratio of the rice line was lower than 1/4, the rice line was classified as resistant (R); when the ratios were between 1/4 and 1, the rice line was rated as susceptible (S). The lesion length 14 days following inoculation was also measured and used to compare with the data obtained 7 days later.

## Data access

Sequence data generated for this study have been submitted to the NCBI Sequence Read Archive under BioProject accession PRJNA350904.

## Acknowledgments

This work was supported by National Key Research and Development Program of China (2017YFD0201100), National Natural Science Foundation of China (NSFC) (31770003, 31970003, 31571974 and 31801697), Science and Technology Department of Hubei Province (2017CFB438).

---

**Figure S1.**
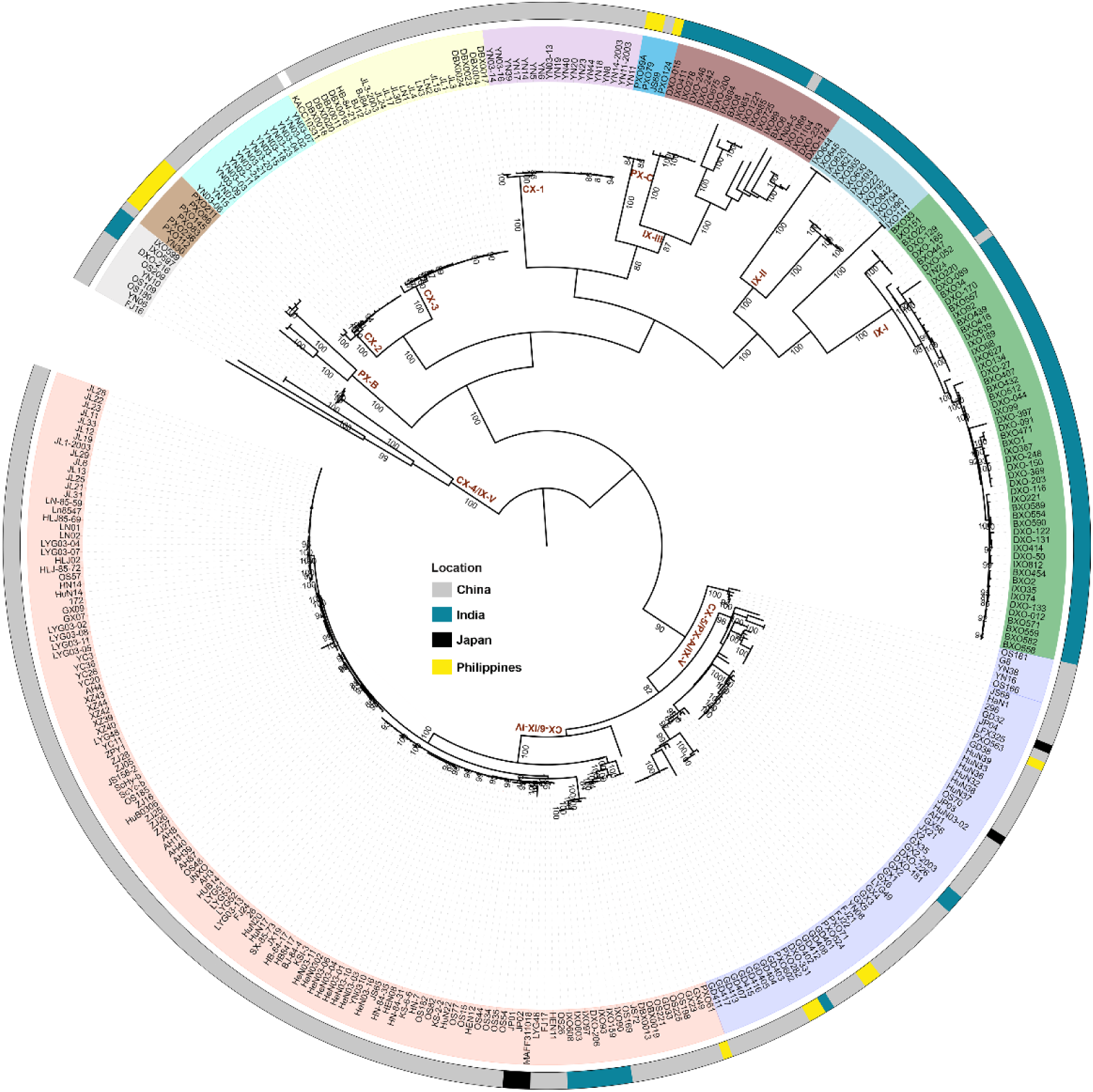
Population structure of Asian *Xanthomonas oryzae* pv. *oryzae*. For phylogenetic analysis, parsnp was used to call SNPs, and RAxML was used to build the ML tree. Background colors of nodes represent different lineages. The outmost circle refers to different countries where Xoo were isolated.

**Figure S2.**
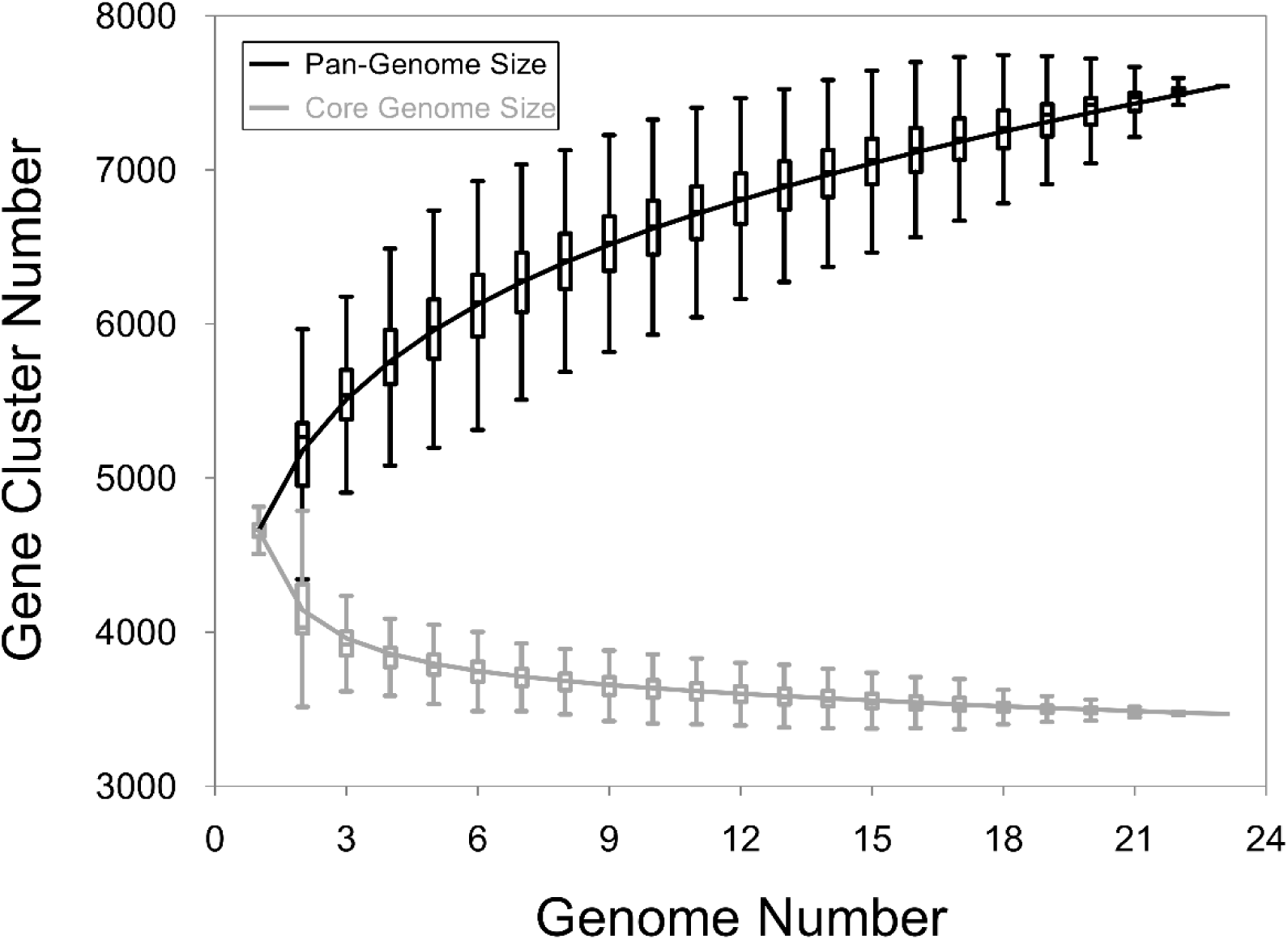
Plot of Pan and core genes from the 22 complete *Xoo* genomes.

**Figure S3.**
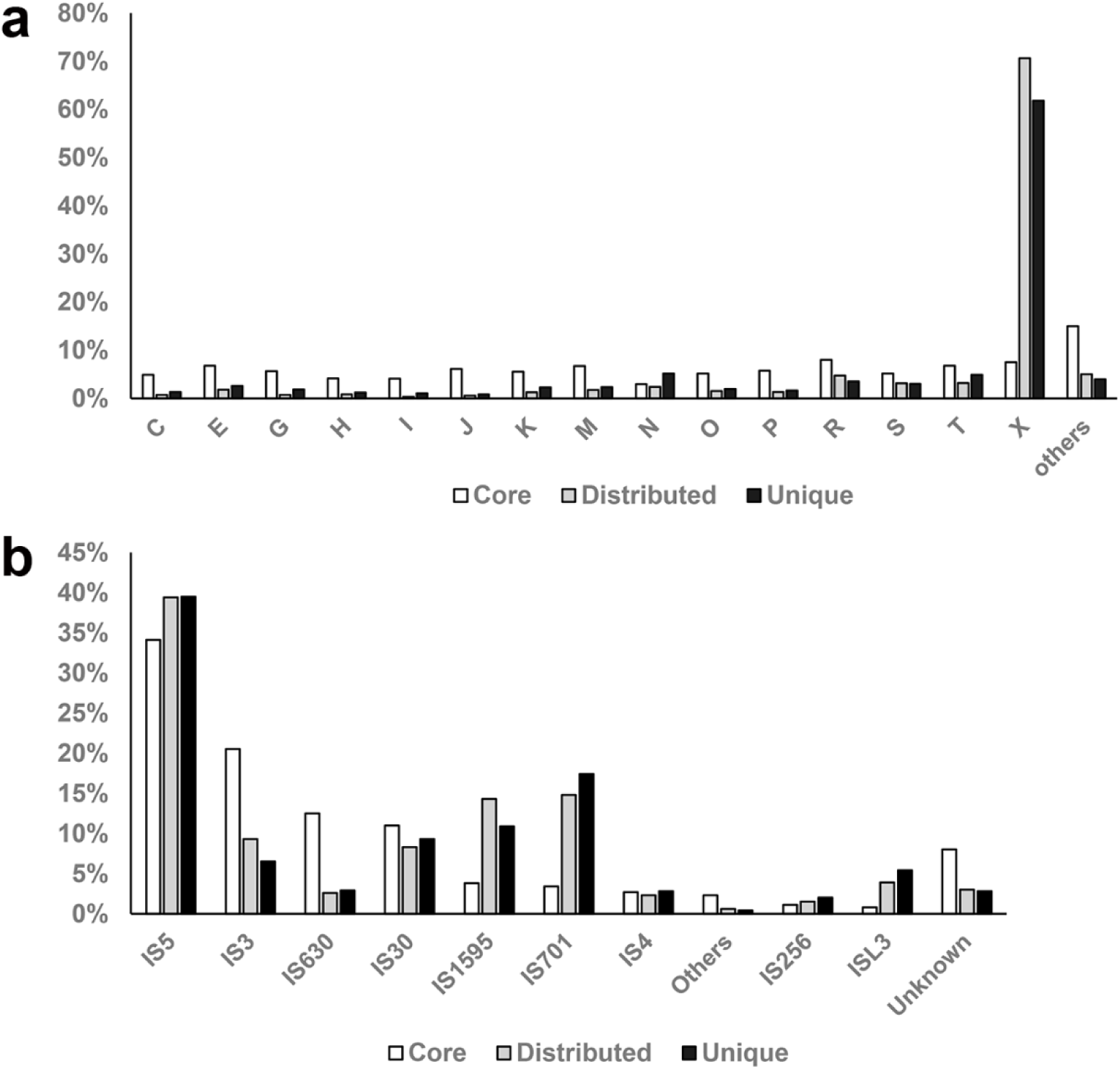
Function analysis of pan genes. (a) COG function of different types of gene families. Core, genes shared by all the 22 genomes; distributed, gene shared by 2 ∼ 21 genomes; unique, genes unique to each genome. (b) Classification of transposase genes. COG categories are energy production and conversion (C), amino acid metabolism and transport (E), carbohydrate metabolism and transport (G), coenzyme metabolism (H), lipid metabolism (I), translation (J), transcription (K), cell wall/membrane/envelop biogenesis (M), cell motility (N), posttranslational modification, protein turnover, and chaperone functions (O), inorganic ion transport and metabolism (P), general functional prediction only (R), function unknown (S), signal transduction (T), Mobilome: prophages, transposons (X).

Table S1 Xoo isolates used in this study.

**Table S2.**
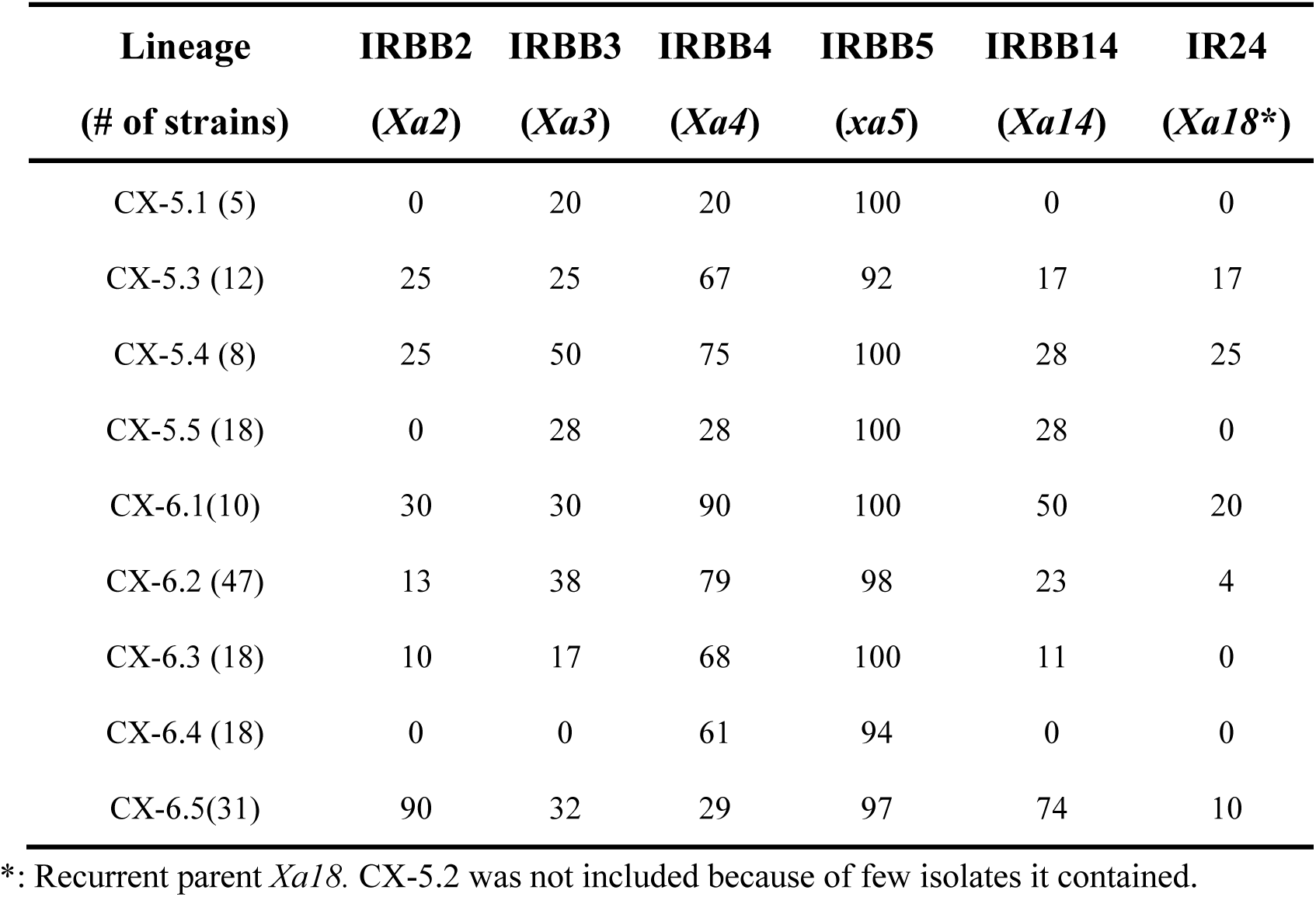
Resistance of different rice lines to *Xoo* strains of different sub-lineages of CX-5 and CX-6. Shown is the percentage of *Xoo* strains of different lineages that are unable to cause disease.

Table S3 Transposase content of Xoo with complete genomes.

Table S4 Distribution of TAL among different *Xoo* with complete genomes

